# Multifactorial and closed head impact traumatic brain injuries cause distinct tactile hypersensitivity profiles

**DOI:** 10.1101/2020.06.01.127944

**Authors:** A-S. Wattiez, W.C. Castonguay, O.J. Gaul, J.S. Waite, C.M. Schmidt, A. Reis, B.J. Rea, L.P. Sowers, C.J. Cintrón-Pérez, E. Vázquez-Rosa, A.A. Pieper, A.F. Russo

## Abstract

Chronic complications of traumatic brain injury (TBI) represent one of the greatest financial burdens and sources of suffering in society today. A substantial number of these patients suffer from post-traumatic headache (PTH), which is typically associated with tactile allodynia. Unfortunately, this phenomenon has been under-studied, in large part due to the lack of well-characterized laboratory animal models. We have addressed this gap in the field by characterizing the tactile sensory profile of two non-penetrating models of PTH. We show that multifactorial TBI, consisting of aspects of impact, acceleration/deceleration, and blast wave exposure, produces long term tactile hypersensitivity and central sensitization, phenotypes reminiscent of PTH in patients, in both cephalic and extracephalic regions. By contrast, closed head injury induces only transient cephalic tactile hypersensitivity, with no extracephalic consequences. Both models show more severe phenotype with repetitive daily injury for three days, compared to either one or three successive injuries in a single day, providing new insight into patterns of injury that may place patients at greater risk of developing PTH. Importantly, even after recovery from transient cephalic tactile hypersensitivity, mice subjected to closed head injury had persistent hypersensitivity to established migraine triggers, including calcitonin gene-related peptide (CGRP) and sodium nitroprusside, a nitric oxide donor. Our results offer new tools for studying PTH, as well as preclinical support for a pathophysiologic role of CGRP in this condition.

**Summary:** Two models of post-traumatic headache after traumatic brain injury provide novel laboratory tools and insights in relative risks of injury and therapeutic opportunities.

## Introduction

Traumatic brain injury (TBI), commonly caused by crashes, falls, contact sports, explosions, or assaults, is a leading cause of chronic morbidity and mortality. Indeed, it is estimated that 69 million people worldwide sustain a TBI every year [14], and in the United States alone there are ∼5 million Americans currently living with TBI-related disability. This translates to an annual cost of ∼$80 billion, with TBI as the leading cause of death in adults under 45 years of age [18; 31; 43]. After the initial physical injury, TBI involves a complex secondary response with prolonged immune reactivity [55], neurodegeneration [46], mitochondrial dysfunction [24], and cerebrovascular compromise [1]. An estimated 70-90% of TBIs are mild [6; 10; 53], defined as suffering from momentary changes in consciousness and scoring between 13 and 15 on the Glasgow Coma Scale [48]. While most patients suffering from mild TBI experience resolution of acute symptoms within 12 weeks, subsequent chronic pathological changes in the brain frequently cause long-term disability [4; 10]. Furthermore, the epidemiologic risk of new TBI is two to five times greater after sustaining a first one [5], such as with repetitive direct impact in athletes or repeated blast exposure in military personnel. Repetitive TBI often leads to severe sensorimotor disability [34], including post-traumatic headache (PTH) associated with severe tactile allodynia [3; 38].

The International Classification of Headache Disorders (ICHD) defines PTH as a secondary headache occurring within 7 days of TBI, or occurring upon regaining of consciousness after trauma [23]. PTH can be acute (resolved within 3 months of onset) or persistent [3], and also shares clinical characteristics of migraine [27; 28]. Despite its high prevalence, however, the pathophysiology of PTH is still poorly understood [3], due in large part to challenges in modeling this condition in animals. Here, we have addressed this gap in the field by characterizing the long-term effects of both single and repeated TBI in mice on a symptom consistent with PTH in people: tactile hypersensitivity in both cephalic and extracephalic regions [29; 33].

Specifically, we compared single and multiple TBIs in: (a) closed head impact to mimic a common form of simple TBI, and (b) multifactorial TBI incorporating concussive impact, acceleration/deceleration, and blast wave exposure to mimic more complex TBI. We asked whether mice after closed head impact were sensitized to pain triggers by using a normally non-noxious dose of two migraine triggers: calcitonin gene-related peptide (CGRP) and the nitric oxide donor sodium nitroprusside (SNP). CGRP and nitric oxide are well-established triggers of migraine in people, and of migraine-like symptoms in rodents [19; 20; 32; 42]. We were particularly interested in whether CGRP might be involved in our mouse models of PTH, given the similarities between PTH and migraine and the recently established efficacy of CGRP-based drugs for migraine prophylaxis and treatment.

## Methods

### Animals

Wildtype outbred CD1 mice were obtained from Charles River Laboratories, Roanoke, IL at 8 weeks of age and acclimated for one week in the animal facility prior to experimental testing. Equal numbers of male and female mice were used. Mice were housed 4 per cage on a 12-h light cycle with food and water ad libitum. All experiments were performed between 8 AM and 3 PM, with mice randomized and investigators blinded to injury status and/or drug treatment. Animal procedures were approved by the University of Iowa Animal Care and Use Committee and performed in accordance with standards set by the National Institutes of Health, policies of the International Association for the Study of Pain and ARRIVE guidelines.

### Drug administration

All injections were performed intraperitoneally (i.p.) with a 0.3 mm × 13 mm needle. Dulbecco PBS (Hyclone) was used as the diluent and vehicle. The amounts injected were as follows: 0.01 or 0.1 mg/kg rat α-CGRP (Sigma-Aldrich, St Louis, MO), 0.25 or 2.5 mg/kg sodium nitroprusside (SNP) (Sigma-Aldrich, St Louis, MO). Animals were gently handled so that no anesthetic agents were needed during injections. All injections were performed by either WCC, BJR, or ASW. For induction of injuries, anesthetics were used: inhaled 5% isoflurane for closed head impact or an i.p. injection of 87.5 mg/kg ketamine, 12.5 mg/kg xylazine for overpressure injury.

### Multifactorial brain injury

Mice were anesthetized with ketamine (87.5 mg/kg) and xylazine (12.5 mg/kg) via i.p. injection, and then placed in an enclosed chamber constructed from an air tank partitioned into two sides and separated by a port covered by a mylar membrane. The pressure in the side not containing the mouse was increased to cause membrane rupture, which generated a 1-2 ms jet airflow blast wave that reached the animal’s head at an average of 25 pounds per square inch. The head remained untethered in a padded holder, while the body was fully shielded by a metal tube. The blast pressure wave produced upon membrane rupture provides a concussive injury, which is followed by acceleration/deceleration of the head and then exposure to the ensuing air wave within an enclosed space. This procedure is routinely performed in our laboratory [17; 51; 57; 58]. Immediately after injury, the mouse was removed from the restraint and placed in its home cage to recover from the anesthetic. Sham mice underwent the same anesthesia, handling and exposure to the noise coming for the rupture of the mylar membrane, but were not placed within the TBI apparatus and therefore not exposed to injury. Mice were monitored until they were fully recovered from anesthesia and able to perform the righting reflex.

### Closed head impact TBI

A previously described model of weight drop-mediated TBI was used [26]. Mice were anaesthetized with 5% isoflurane for ∼2 min, then immediately placed on a foam cushion in a head trauma device. The instrument consists of a fiberglass tube (80 cm high, inner diameter 13 mm), placed vertically over and lightly touching the head. A 30 g metal weight was then dropped from the top of the tube to strike the head at the temporal right side between the corner of the eye and the ear. The foam cushion prevented rotation of the head. Immediately after impact, mice were placed in their home cage and observed until their righting reflex returned. Sham-treated animals were anaesthetized but not subjected to the weight drop.

There were no visible physical signs of trauma in any of the animals exposed to either form of TBI, and all animals recovered similarly to sham mice. In each injury group, 2 animals (1 male and 1 female) were euthanized 5 min after recovery from the last injury and the skull was exposed. There were no signs of skull fracture or brain hemorrhage in any injury paradigm. Therefore, the injuries are characterized as mild.

### Periorbital and plantar tactile sensitivity

For periorbital testing, mice were tested as described by [9]. Briefly, each mouse was acclimated to its own polycoated paper cup (Choice 4 oz. paper cups; 6.5 cm top diameter, 4.5 cm bottom diameter, 72.5 cm length) for 20 min each day for 5 to 10 days, until habituated. During each acclimation period, a von Frey filament was repeatedly approached to the periorbital area without applying pressure, in order to decrease their withdrawal reflex. Mice were considered habituated once the tip of the filament could reach the head (without any pressure) without the mouse reacting to it.

For plantar testing, mice were habituated to an acrylic chamber (dimensions 114 x 80 cm) placed over a grid support (Bioseb, France), for 2 h on the day before the first testing, and for 30 min immediately before testing.

A set of eight von Frey filaments was used from A (0.008 g) to H (1 g) (Bioseb, France). Testing was performed according to the up and down method previously described [11; 16]. Briefly, filaments were applied for 3 s at the periorbital area or 5 s at the plantar area. The D (0.07 g) filament was applied first. If the animal reacted (withdrew the head, wiped eyes or periorbital region, withdrew, shook, or licked hindpaw for plantar), then a lower filament was applied. If there was no reaction, then a higher filament was applied. A pattern was recorded. This method was used until 5 applications after a first change in the pattern were assessed. The last filament applied and the final pattern were used to calculate the 50% threshold following an established equation [11; 16]. Since this technique does not yield continuous thresholds and data cannot be analyzed using parametric statistics, the 50% thresholds (g) were log-transformed before being analyzed in order to obtain normally distributed data.

### Experimental design

In order to reduce the number of animals used in this study, the same cohorts of mice were tested for periorbital and plantar hypersensitivity after respective injuries (multifactorial injury in Fig. 1A and 1B, and closed head impact injury in Fig. 2A and 2B). However, because we wanted to limit the number of time mice were injected with treatments within the same week. different cohorts of mice were tested for periorbital and plantar hypersensitivity after sub-threshold triggers in Fig. 3, 5B and 5C (periorbital) and 4, 5D and 5E (plantar). In Fig.4B, one of the cohorts tested on D21 generated aberrant results and exhibited agitated behavior all through assay. For this reason, the testing was repeated on D22, when their behavior appeared normal again. The results obtained on D22 were used for the present results, and the aberrant data of D21 were discarded.

**Figure 1.**
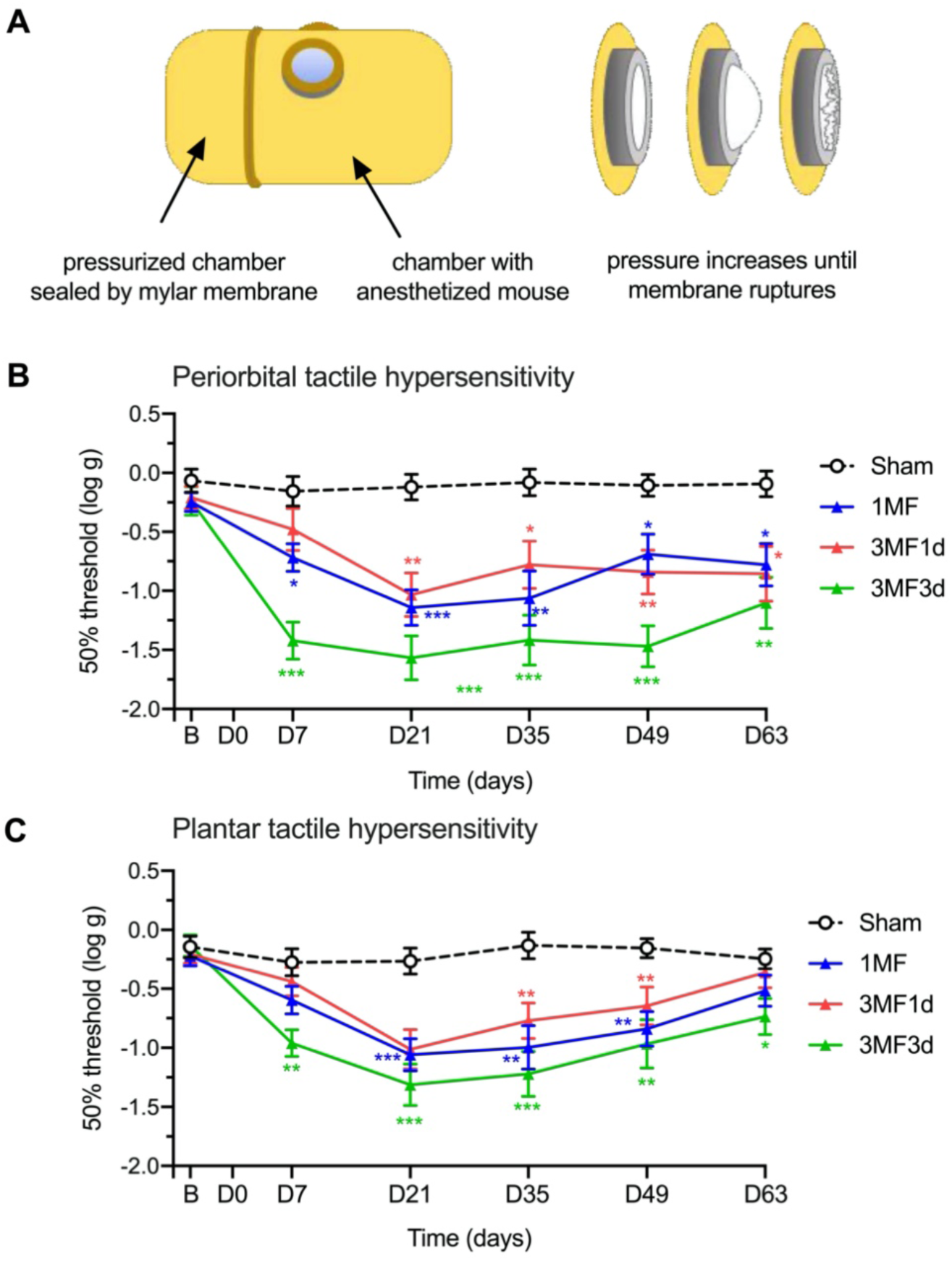
Multifactorial TBI induces persistent tactile hypersensitivity. **A.** Schematic of the overpressure chamber in which multifactorial injury was induced. Anesthetized mice were placed in an enclosed chamber partitioned into two sides, sealed by a mylar membrane. Pressure was gradually increased in the portion of the chamber without the mouse until the membrane ruptured, generating an overpressure air wave reaching the animal’s head at a pressure of ∼25 PSI. **B.** Periorbital tactile hypersensitivity was assessed before injury at baseline (B), and at days 7, 21, 35, 49, and 63 after either a sham injury (n=19), a single multifactorial (MF) injury (1MF, n=20), 3 injuries within the same day (3MF1d, n=17), or 3 injuries over 3 consecutive days (3MF3d, n=14). **C.** Plantar tactile hypersensitivity on the same mice as in panel B. The mean ± SEM 50% thresholds are presented. Statistics are described in Table 1.

**Figure 2:**
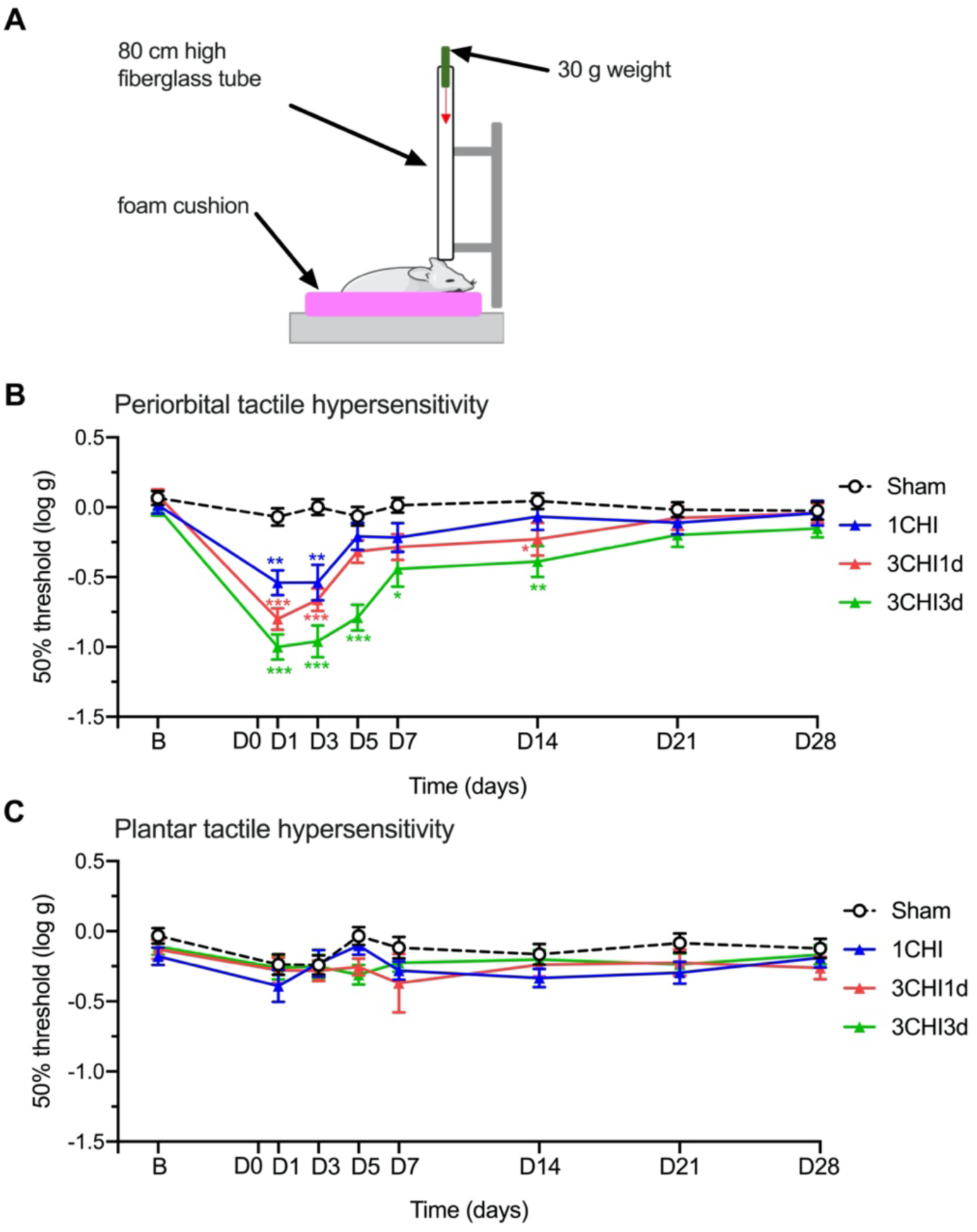
Closed head impact TBI induces transient periorbital tactile hypersensitivity but not plantar tactile hypersensitivity. **A.** Schematic of the weight-drop apparatus in which the closed head impact injury was induced. Anesthetized mice were placed under a fiberglass tube (inner diameter 13 mm), placed vertically over the mouse head. A 30 g metal weight was then dropped from the top of the tube (80 cm) to strike the head at the temporal right side between the corner of the eye and the ear. A foam cushion was placed underneath the animal to protect it from contact with the surface beneath the head trauma device and to prevent any rotational damage of the head. **B.** Periorbital tactile hypersensitivity was assessed at baseline (B) before injury, at days 1, 3, 5, 7, 14, 21, and 28 after either a sham injury (n=16-18), a single closed head impact injury (1CHI, n=14-16), 3 injuries within the same day (3CHI1d, n=16-17), or 3 injuries over 3 consecutive days (3CHI3d, n=18). **C.** Plantar tactile hypersensitivity on the same mice as in panel B. The mean ± SEM 50% threshold are presented. Statistics are described in Table 1.

**Figure 3:**
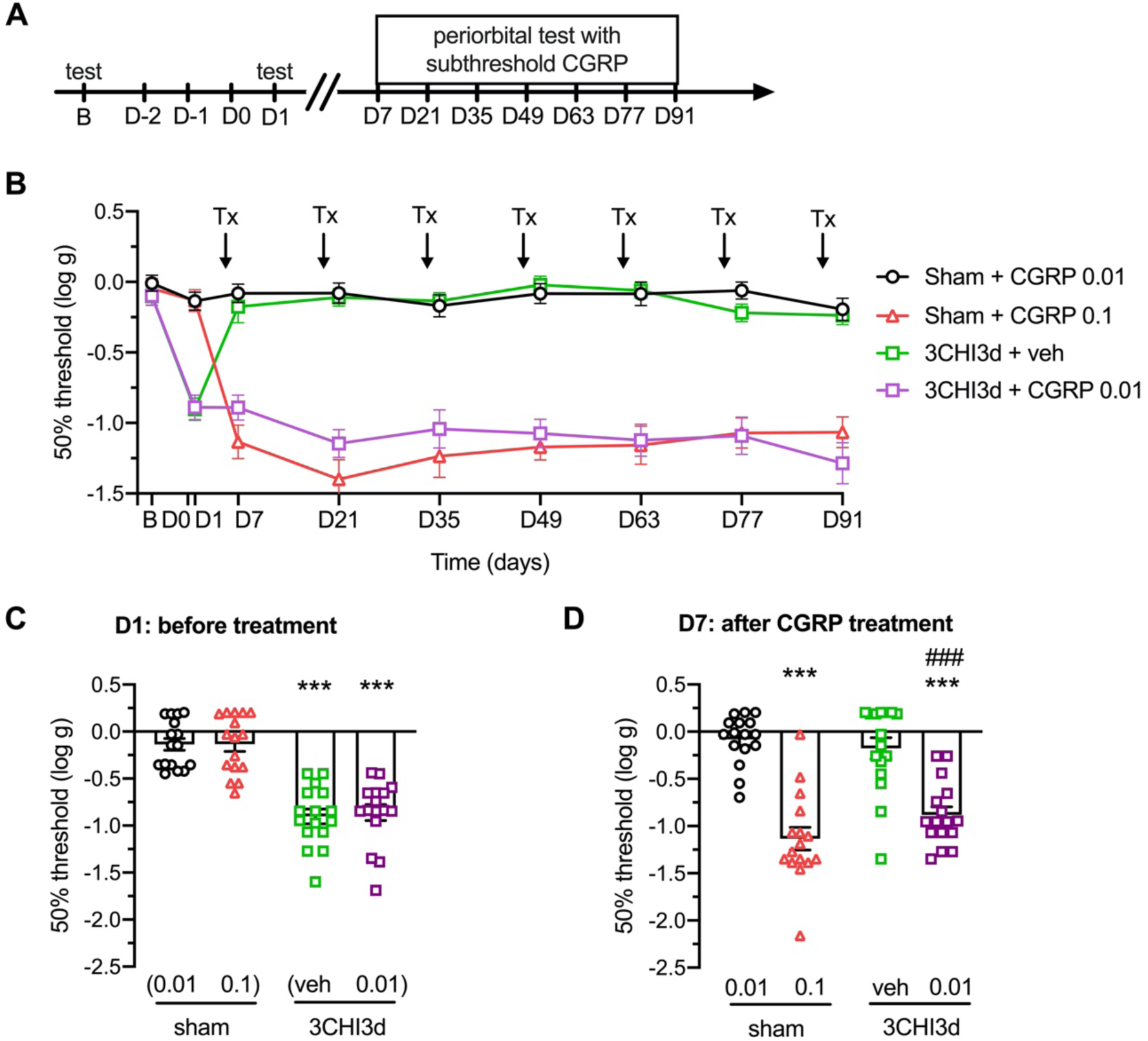
Closed head impact TBI sensitizes mice to non-noxious CGRP in the cephalic area. **A.** Schematics of the experimental procedure. **B.** Periorbital tactile hypersensitivity was assessed before sham or injuries (B), one day after the last sham or closed head impact injury (3CHI3d model) without any treatment (D1), and then 30 min after administration of vehicle, non-noxious dose of CGRP (0.01 mg/kg, i.p.), or normal dose of CGRP (0.1 mg/kg, i.p.) at days 7, 21, 35, 49, 63, 77, and 91 after the last sham/injury. **C.** Scatter plot representation of the individual thresholds for mice at D1 of the experiment presented in B. At D1, mice are 1 day post sham or injuries, and did not receive any treatment. **D.** Scatter plot representation of the individual thresholds for mice at D7 of the experiment presented in B. At D7, mice are 7 days post sham or injuries, and were tested 30 min after administration of treatments. The mean ± SEM 50% threshold are presented in all graphs. N=15-16 per group. Statistics are described in Table 1.

**Figure 4:**
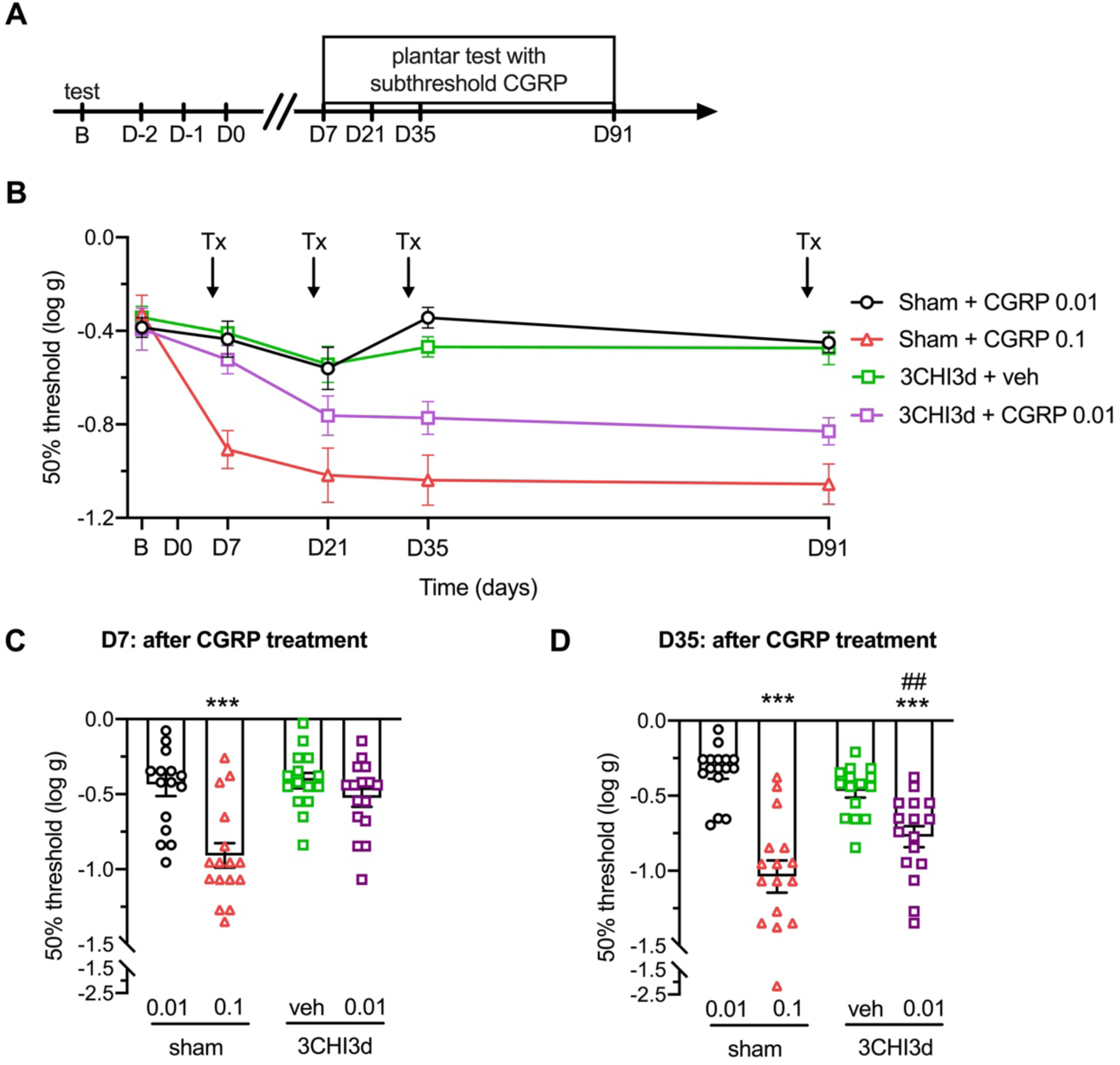
Closed head impact TBI sensitizes mice to non-noxious CGRP in the extracephalic area. **A.** Schematic of the experimental procedure. **B.** Plantar tactile hypersensitivity was assessed before sham or injuries (B), one day after the last sham or closed head impact injury (3CHI3d model) without any treatment (D1), and then 30 min after administration of vehicle, non-noxious dose of CGRP (0.01 mg/kg, i.p.), or normal dose of CGRP (0.1 mg/kg, i.p.) at days 7, 21, 35, and 91 after the last sham/injury. **C.** Scatter plot representation of the individual thresholds for mice at D7 of the experiment presented in B. At D7, mice are 7 day post sham or injuries, and were tested 30 min after administration of treatments. **D.** Scatter plot representation of the individual thresholds for mice at D35 of the experiment presented in B. At D35, mice are 35 days post sham or injuries, and were tested 30 min after administration of treatments. The mean ± SEM 50% threshold are presented in all graphs. N=15-16 per group. Statistics are described in Table 1.

**Figure 5:**
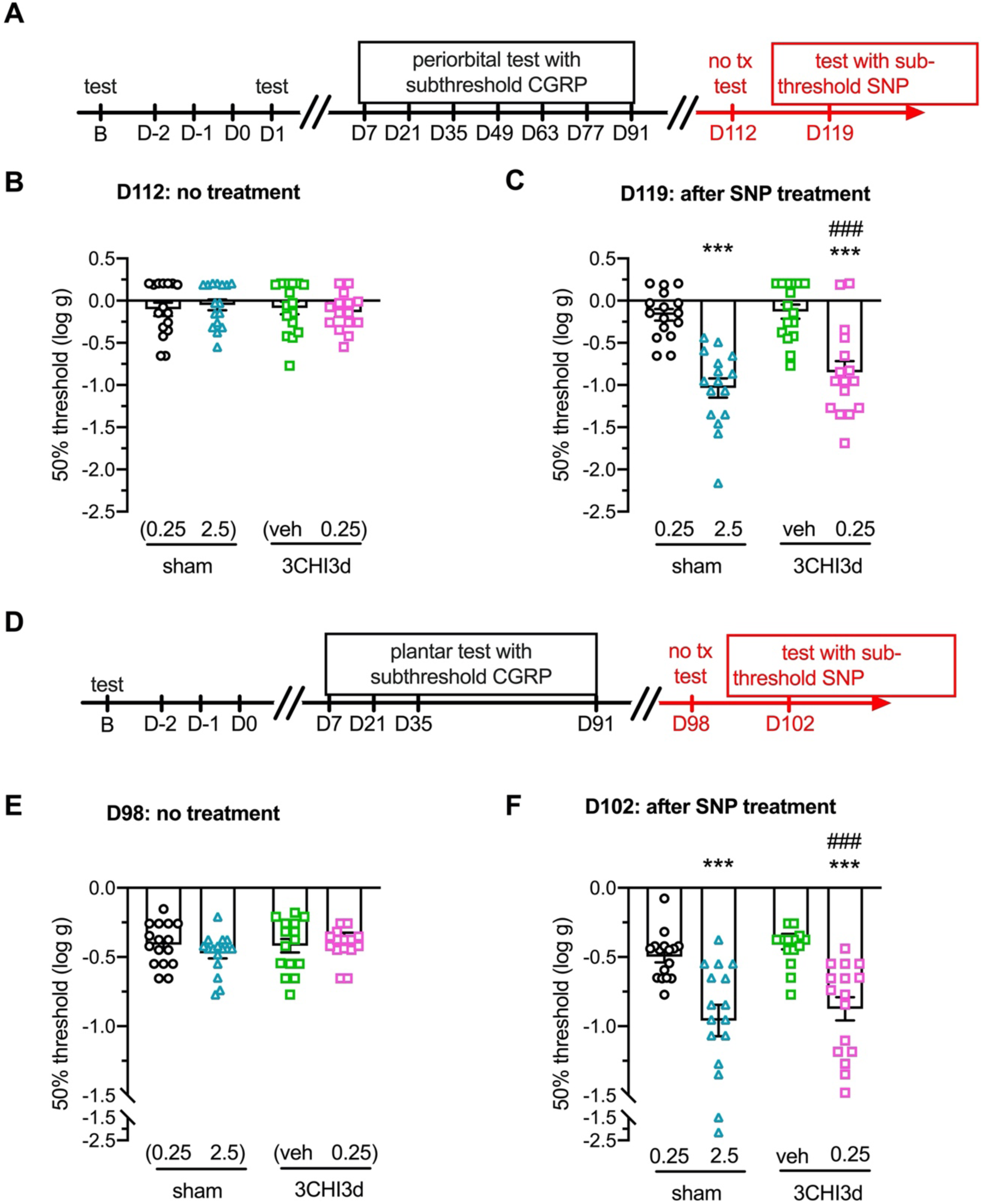
Closed head impact TBI sensitizes mice to non-noxious SNP in cephalic and extracephalic areas. **A.** Schematic of the experimental procedure for B and C. This experiment uses the same animals as in Fig. 3 at later time-points (highlighted in red). Groups were composed of the same animals, only the treatments were changed. **B.** Scatter plot representation of the individual thresholds for mice at D112 of the experiment presented in Fig. 3B. At D112, mice are 112 days post sham or injuries, and were tested without any treatment. **C.** Scatter plot representation of the individual thresholds for mice at D119 of the experiment presented in Fig. 3B. At D119, mice are 119 days post sham or injuries, and were tested 60 min after administration of either vehicle, non-noxious dose of SNP (0.25 mg/kg, i.p.), or normal dose of SNP (2.5 mg/kg, i.p.). **D.** Schematic of the experimental procedure for E and F. This experiment uses the same animals as in Fig. 4 at later time-points (highlighted in red). Groups were composed of the same animals, only the treatments were changed. **E.** Scatter plot representation of the individual thresholds for mice at D98 of the experiment presented in Fig. 4B. At D98, mice are 98 days post sham or injuries, and were tested without any treatment. **F.** Scatter plot representation of the individual thresholds for mice at D119 of the experiment presented in Fig. 4B. At D102, mice are 102 days post sham or injuries, and were tested 60 min after administration of either vehicle, non-noxious dose of SNP (0.25 mg/kg, i.p.), or normal dose of SNP (2.5 mg/kg, i.p.). The mean ± SEM 50% threshold are presented in all graphs. N=15-16 per group. Statistics are described in Table 1.

For each experiment, mice were brought to the behavioral room for acclimation for one hour prior to handling/testing. Baseline 50% thresholds were recorded for each animal before TBI. Animals were then separated into experimental groups using a block randomization protocol per cage (all 4 animals within a same cage were allocated to the same injury group in case they develop anxiety, and in order to not stress sham animals). Animals in sham groups received anesthetics on 3 consecutive days in order to provide the most robust negative control. All subsequent time-points are expressed as days after the last sham or injury exposure. When treatments were administered, animals were placed back into their home cages immediately after drug administration and before the beginning of testing. Mice were tested 30 min after CGRP and 1 h after SNP administration.

**Table 1:**
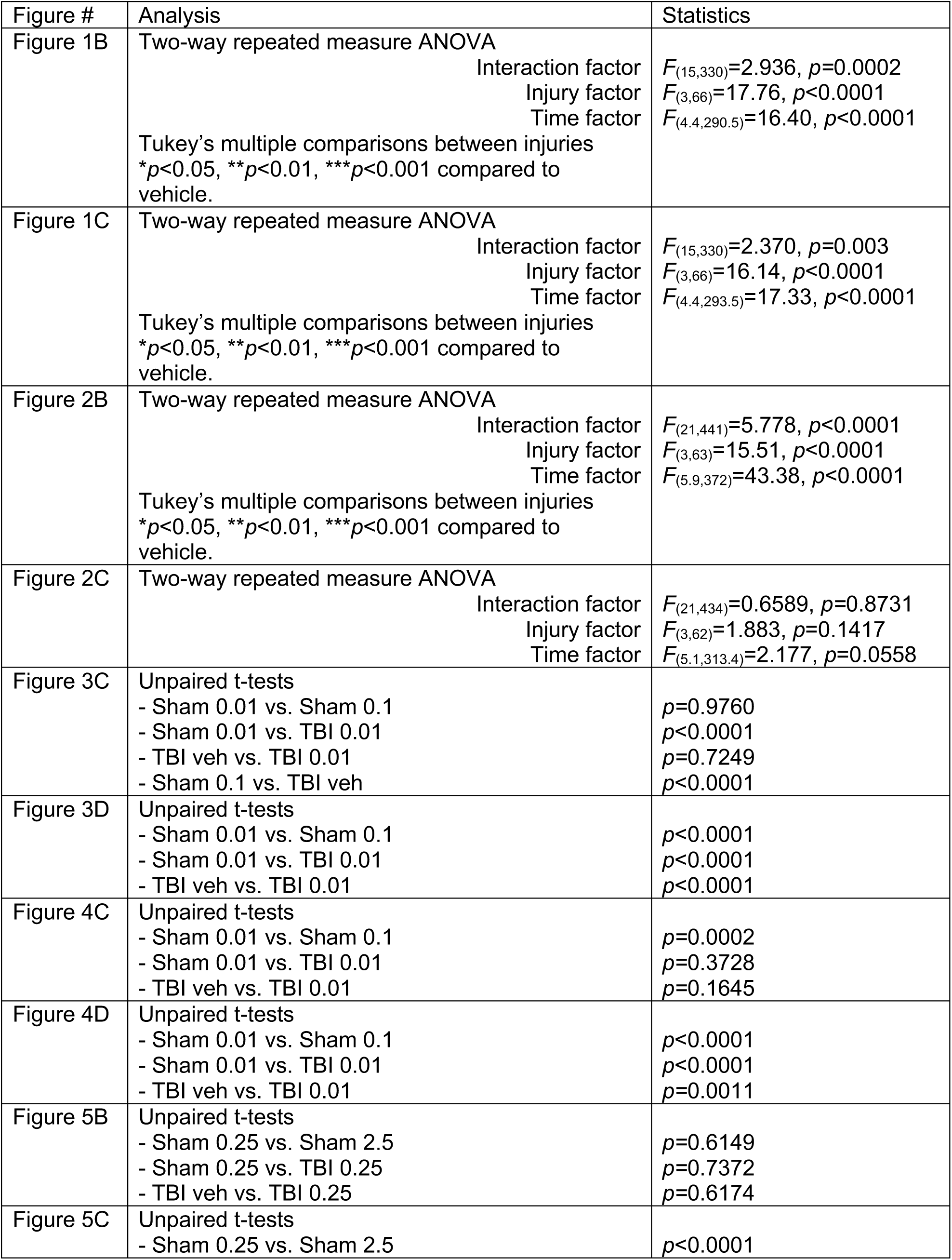
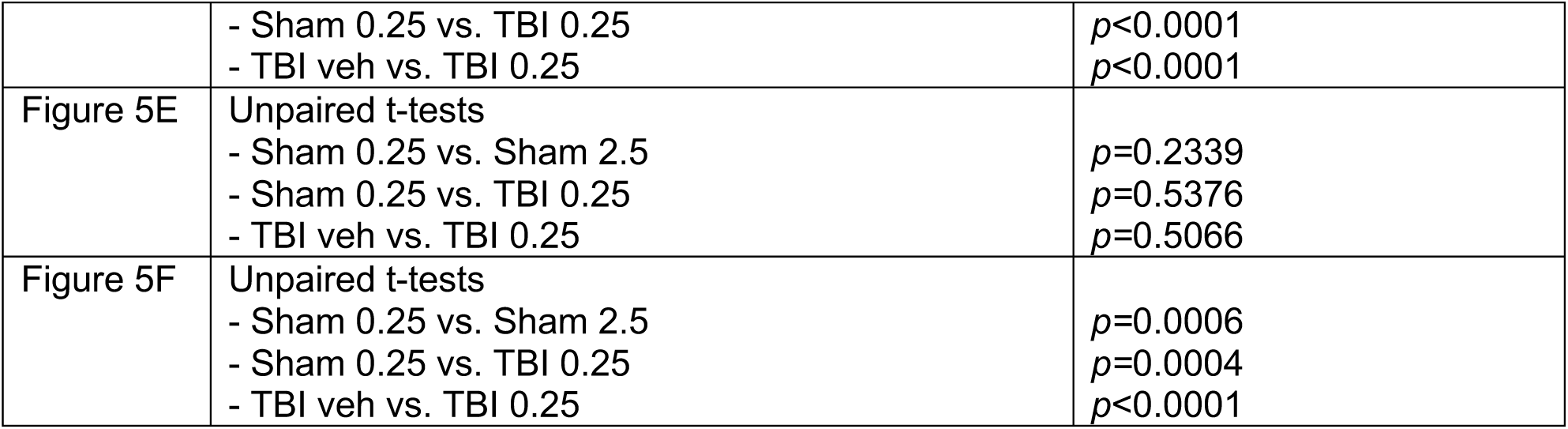
Statistical analyses.

### Statistical analysis

A power analysis was performed for sample size estimation using ClinCalc.com. The effect size of this study was estimated at 50% decrease for both tactile hypersensitivity and light aversion, based on data from previous similar studies. With an alpha of 0.05 and power at 0.80, the projected sample size needed with this effect size was approximately 14 per group for tactile hypersensitivity and 15 for light aversion. Data are reported as mean ± SEM. Data were analyzed using GraphPad Prism 8.4 software (RRID: SCR_002798). Significance was set at *p*<0.05. No sex-specific differences were observed in this study, and the effect size was powered for analysis of males and females pooled together.

All statistics are reported in Table 1. For Fig. 1 and 2, data are plotted as a function of time (line graphs), and a two-way repeated measure ANOVA was performed (factors time and injury). For all graphs, the interaction between the two factors was significant, therefore a Tukey multiple-comparison test was performed to compare the effect of each injury at each time point. For Fig. 3B and 4B, no statistics were performed because not all injury groups received all treatments. Instead, at the different time points, statistics were assessed and presented on the scatter plots in Fig. 3C, 3D, 4C, 4D, 5B, 5C, 5E, 5F. For the scatter plots, since a two-way ANOVA was not possible, unpaired t-tests were used to assess the difference between 2 injury/treatment groups.

Exclusions were applied to the datasets for the following reasons: fight wounds that would have affected pain thresholds (n=3) or development of dermatitis that would have affected pain threshold (n=1). All data from those animals was excluded from the study. Additionally, over the 58 animals receiving the multifactorial TBI (134 injuries performed overall), 5 died on injury day (4 females, 1 male). Those animals were alive at the end of injuries but never recovered from anesthesia. There was no animal loss during the closed head impact procedure.

## Results

### Multifactorial brain injury induces persistent tactile hypersensitivity

The murine multifactorial brain injury model incorporates concussion and acceleration/deceleration upon exposure to an overpressure air wave, resulting in neurodegeneration and neurobehavioral deficits reminiscent of multifactorial TBI in people [17; 51; 57; 58]. Exposure of mice to this injury (Fig. 1A) decreased the 50% thresholds (log transformed to obtain normal distribution) in the cephalic region (periorbital, on the midline just above the eyes) starting 7 days after injury and lasting for at least 9 weeks (D63, Fig. 1B). All three groups, single multifactorial injury (1MF), 3 multifactorial injuries within the same day (3MF1d), and 1 multifactorial injury every day for 3 consecutive days (3MF3d), significantly decreased thresholds compared to the sham injury group. The most severe effect was observed in 3MF3d mice (Fig. 1B).

We also evaluated tactile sensitivity in the plantar area of the paw to assess for central sensitization, which is associated with development of chronic pain. Similar to what was observed in the cephalic area, all groups of multifactorial TBI showed significantly decreased plantar thresholds compared to the sham injury group, from 3 weeks (D35) after injuries and for at least 7 weeks (D49, Fig. 1C). Once again, the most robust phenotype was observed in the 3MF3d group, with the decrease in thresholds reaching significance more rapidly (1 week (D7) after injury) and lasting longer (at least 9 weeks (D63)) (Fig. 1C). There was no difference observed between males and females in periorbital or plantar thresholds before or after injury in any of the groups (data not shown).

### Closed head impact TBI induces transient periorbital but not plantar tactile hypersensitivity

We next assessed the same phenotypes in a closed head impact model of TBI induced by dropping a weight onto the intact skull (Fig. 2A). The same single and multiple injury schedule was used as with the multifactorial TBI model. All three groups, single closed head impact injury (1CHI), 3 injuries within 1 day (3CHI1d), or 1 injury a day for 3 consecutive days (3CHI3d), produced a decrease in cephalic 50% thresholds within 1 day (Fig. 2B). However, in contrast to multifactorial TBI, this effect was transient in all groups. The peak decrease was observed at 1 day after injury, and followed by slow recovery to sham levels at day 5 in 1CHI and 3CHI1d groups. As with multifactorial TBI, the most robust effect was observed in the multiple injuries over 3 days group, for which thresholds took longer (1 to 2 weeks) to return to normal (sham) levels (Fig. 2B). However, in contrast to what was observed with multifactorial TBI, no form of closed head impact induced a change in plantar thresholds (Fig. 2C). No differences were observed between males and females in this model.

### Closed head impact sensitizes mice to non-noxious CGRP in the cephalic area

Considering that the closed head impact TBI model produced only transient symptoms, we decided to investigate whether these mice were sensitized to pain triggers around the time of recovery from periorbital tactile hypersensitivity. To begin, we tested a normally non-noxious dose of CGRP. CGRP is known to be involved in migraine, and peripheral administration of CGRP to mice induces migraine-like light aversion [32] and facial grimace that has been established to indicate the experience of spontaneous pain [44]. Here, we observed that the CGRP dose classically used to elicit light aversion and grimace (0.1 mg/kg i.p.) also caused periorbital tactile sensitivity in sham mice (Fig. 3B, D). By contrast, sham mice given a 10-fold lower dose (0.01 mg/kg i.p.) were unaffected (Fig. 3B, D), and thus this was defined as the non-noxious dose.

The 3CHI3d mice were then given either vehicle or a subthreshold, non-noxious dose of CGRP as a subsequent trigger one week after injury. The 3CHI3d vehicle mice given vehicle displayed only a transient decrease in withdrawal thresholds before vehicle was administered, and thresholds were no different from baseline after each vehicle administration (Fig. 3B, C). However, the 3CHI3d mice treated with subthreshold CGRP one week after injury displayed a robust and significant decrease in thresholds, compared to both the sham group receiving the same dose, and the 3CHI3d group receiving vehicle (Fig. 3B, D). This sensitization persisted until at least day 91 (Fig. 3D).

### Closed head impact sensitizes mice to non-noxious CGRP in the extracephalic area

We next tested whether the same non-noxious dose of CGRP could also decrease tactile thresholds on the paw after multiple closed head impact injuries, as it did on the head (Fig. 4A details the experimental procedure). As with periorbital sensitivity, i.p. injection of CGRP at 0.1 mg/kg caused plantar tactile sensitivity in sham mice (Fig. 4B, D). In contrast, sham mice given a 10-fold lower dose (0.01 mg/kg i.p.) were unaffected (Fig. 4B, D), so this dose was considered non-noxious for extracephalic testing just as it was for periorbital testing.

As seen with the periorbital area, 3CHI3d mice that received this non-noxious dose of CGRP showed a significant decrease in 50% thresholds compared to vehicle and sham mice receiving the same CGRP dose (Fig. 4B). However, the effect was somewhat less robust than seen with the periorbital region. Specifically, it was not apparent one week after injury (Fig. 4B, C) and was only detected 3 weeks after the last injury (Fig. 4B, D). This effect persisted until at least day 91.

### Closed head impact sensitizes mice to non-noxious sodium nitroprusside in both the cephalic and extracephalic areas

We next assessed the effect of a non-noxious dose of another migraine trigger, the nitric oxide donor sodium nitroprusside (SNP) [19] (Fig. 5A). Like CGRP, nitric oxide donors can be used pre-clinically to induce migraine-like phenotypes in mice and rats [42]. To compare with CGRP, we used the same mice used for the subthreshold experiments. Thus, the first step was to determine whether previously administered subthreshold CGRP caused a permanent sensitization of 3CHI3d mice. We re-assessed the cephalic threshold without administering vehicle or CGRP (see Fig. 5A for experimental procedure) and observed that thresholds for all groups at day 112 were back to baseline (Fig. 5B). This indicates that the evoked pain induced by subthreshold CGRP is not permanent.

As with CGRP, we observed that SNP (2.5 mg/kg i.p.) induced tactile hypersensitivity in sham mice (Fig. 5C). By contrast, mice given a 10-fold lower dose of SNP (0.25 mg/kg i.p.) were unaffected (Fig. 5C). This dose was therefore considered to be non-noxious in this assay. While the non-noxious dose did not induce any threshold change in sham animals, it did significantly decrease thresholds of 3CHI3d mice at 119 days post-injury (Fig. 5C). This decrease was significant compared to both sham mice given 0.25 mg/kg SNP and 3CHI3d mice given vehicle.

These experiments were then repeated for the plantar cohorts (see Fig. 5D for experimental procedure). Ninety-eight days after the last injury, mice were tested without treatment and the plantar thresholds were observed to be back to baseline (Fig. 5E). Thus, as with the periorbital region, trigger-induced plantar tactile hypersensitivity was not permanent. As with periorbital sensitivity, at day 102 after injury, 3CHI3d mice also showed plantar sensitization to the non-noxious dose of SNP (Fig. 5F).

## Discussion

To our knowledge, this is the first direct comparison of chronic nociceptive behaviors related to PTH induced by 2 different models of repetitive TBI. Here, we report that multifactorial TBI and closed head impact TBI induce unique patterns of tactile hypersensitivity. In multifactorial TBI, tactile hypersensitivity develops early in both cephalic and extracephalic areas and persists for at least 9 weeks. By contrast, only cephalic tactile hypersensitivity develops in closed head impact TBI, and this is transient, lasting between 1 and 7 days after injury. After that point, mice display prolonged tactile hypersensitivity in response to normally sub-threshold doses of migraine triggers.

### Choice of TBI models

An increasing number of animal models for TBI are being developed, each with its own unique strengths and weaknesses [56]. The field has historically applied different models considered uniquely appropriate for different studies. Here, we assessed tactile hypersensitivity over time after TBI, a common symptom present in more than half of mild TBI patients that can be attributed to PTH and/or generalized pain [3; 39; 41]. The occurrence of this symptom seems to be independent of the etiology of the injury, although to our knowledge this has not been rigorously investigated. We chose two translatable non-penetrating models with mechanisms close to human TBI injury biomechanics that produce mixed diffuse-focal injuries from a focal impact moving the brain in the skull [48]. Although there are no criteria in animal models to characterize an injury as mild, moderate or severe, the lack of visible damage (no skull fracture or brain hemorrhage) and the prompt recovery of animals seem to indicate that the sustained injuries were relatively mild [15]. Moreover, there were no deaths in the closed head impact group. However, 4 females and 1 male did not recover from anesthesia after multifactorial injury, while all sham injury animals survived, indicating that multifactorial TBI, possibly in combination with anesthetics, may be somewhat more severe. A contribution from anesthetics is suggested by a report of lower mortality after TBI in awake compared to anesthetized animals [50]. The higher ratio of female mice dying from multifactorial injury might be attributed to sex-specific factors, as multiple studies show that while men are more likely to sustain a TBI, women are at greater risk for poor outcomes [45]. Furthermore, a recent finding in rats reported sexual dimorphism, with females displaying a prolonged state of cephalic hyperalgesia [8]. Aside from mortality, we did not detect any sex-specific outcomes in any of our models of TBI, although the studies were not powered to specifically address this question.

### Long-term vs transient hypersensitivity induced by TBI

We found that multifactorial brain injury induced persistent periorbital and plantar tactile hypersensitivity in mice. This result is consistent with a previous study showing persistent (8 weeks) whisker-pad tactile hypersensitivity after a single direct bullet-less gun-induced TBI in rats estimated at >70 PSI [49]. Interestingly, in a subsequent study, this phenotype was not produced when the TBI was performed in awake animals [50], despite the known protective effect of anesthetics [30]. It was hypothesized that the anesthetized model may have a more robust blast wave dissipation due to better placement of the apparatus on the animal, which would result in a more severe injury [50]. The same team also showed that a single injury induced persistent spontaneous pain, as characterized by the rat grimace scale [49; 50]. By comparison, we observed that closed head impact in mice induced transient periorbital tactile hypersensitivity, consistent with previous studies in both mice and rats [7; 8; 37; 40]. However, we did not observe any plantar hypersensitivity. Plantar hypersensitivity is controversial, with some studies reporting sensitivity in rats [7; 8] and others reporting only a transient plantar hypersensitivity in C57BL6/J mice [37; 40]. As our experiments were conducted in CD1 mice, the difference of mouse strains may be a possible explanation for the discrepancy.

### Single vs. repeated TBI

The present findings show in both models that the more injuries received, the more robust the PTH-like phenotype. In patients, it is increasingly reported that subsequent brain injuries are associated with more severe pathology and symptomatology, and it is widely presumed that morbidity after repeated TBI will be proportional to the number of brain injuries sustained [5; 12; 34]. In a review, Hall and colleagues point out that only one study using repeated weight-drop injury models showed a scaled response with increasing number of TBI [5; 47]. All other studies, including ours, show a cumulative but not a strictly additive effect.

With some injuries going unreported [35], athletes may sustain multiple injuries within the same day, or have long inter-injury intervals. For this reason, we chose to compare different injury paradigms. The preclinical literature is inconsistent on the topic, but the majority of studies seem to indicate that inter-injury intervals of less than 5 days seem to worsen the symptoms [36; 52], while injuries sustained months apart seem to either ameliorate [22; 36] or not change [36] the outcome. Interestingly, a pattern of 3 injuries received over 3 consecutive days was more deleterious in our models than 3 injuries within the same day. However, a caveat of our study lies with the likely neuroprotective effect of anesthetic. Animals injured 3 times within the same day had to be sequentially re-anesthetized three times, which could have affected the depth of anesthesia and minimized damage. As with all preclinical studies, the relevance of these results to the human condition in which patients are not anesthetized before TBI must be interpreted with caution. Alternatively, there could be effects of combining a new primary injury with a developing secondary injury. At this time, it is difficult to choose the optimal interval between injuries in an animal model.

### Mild closed head impact sensitizes to sub-threshold migraine triggers after symptoms resolution

Patients who recover from TBI-induced symptoms within the first few weeks/months after injury often develop chronic neurologic changes with sensitization to stressors [13]. Hence, we tested whether TBI sensitized the mice to subthreshold doses of CGRP. Given that persistent hypersensitivity was observed in the multifactorial TBI model, we used the closed head impact mice to address this question. We chose CGRP as our initial stressor because clinical similarities between TBI-induced PTH and primary headaches suggests that PTH could be triggered through the same signaling pathways [3]. The neuropeptide CGRP is also recognized as a critical player in the pathophysiology of migraine [54]. In mice, at a dose of 0.1 mg/kg i.p., CGRP induces spontaneous pain [44] and light-aversion [32]. In this study, we have extended these symptoms to include both periorbital and plantar tactile hypersensitivity. While a 10-fold lower dose is normally unable to induce those symptoms in mice, we show that it does provoke both periorbital and plantar tactile hypersensitivity in mice after repeated injuries. Remarkably, this phenotype lasted for more than 100 days after injury. This sensitivity to CGRP after TBI is reminiscent of the increased sensitivity to CGRP of migraine patients [2; 20; 21; 25].

Similarly, closed head impact mice can also re-develop tactile periorbital and plantar sensitivity after administration of subthreshold SNP (10-fold lower dose than normal), another migraine trigger [19]. This is consistent with previous findings of tactile hypersensitivity after administration of a low dose of nitroglycerin to rats and mice [7; 8; 37]. Both SNP and nitroglycerin are nitric oxide donors that induce migraine-like symptoms in part by recruiting CGRP [3]. An additional piece of evidence highlighting the similarities between migraine and PTH in animal models is that after closed head impact, mice become sensitized and develop periorbital hypersensitivity when exposed to bright light [40]. Those results raise the possibility that infusion of CGRP or SNP in TBI patients might trigger migraine-like headache like they do in migraineurs. This also further supports the likely possibility that TBI patients could respond to CGRP targeting drugs, such as CGRP and receptor blocking antibodies, or small molecule antagonists (gepants), to prevent or treat headache. A few preclinical studies support this idea, although closed head impact was the only model tested for this hypothesis [7; 8; 37; 40]. Indeed, the first clinical trial is ongoing now to evaluate efficacy and tolerability of the CGRP receptor antibody erenumab for treatment of patients suffering from persistent PTH (NCT03974360).

### Conclusion

Multiple animal models are needed to investigate the PTH-like symptoms experienced by patients suffering from the wide variety of TBIs. Here, we have characterized a multifactorial model that mimics persistent hypersensitivity and central sensitization, and a closed head impact model that mimics persistent sensitization to subsequent migraine triggers. The ability to model sustained and transient symptoms is reminiscent of the variability in persistence and manifestation of headache and pain in TBI patients [4; 5; 27; 41]. Furthermore, our results with these models offer support for a causative role of CGRP in PTH, underscoring the potential utility of CGRP-targeting drugs for treatment of TBI-induced PTH and pain symptoms.

## Acknowledgments

The authors would like to thank Dr. Dan Levy and Dr. Dara Bree for teaching the closed head impact model to A-S.W. and L.P.S.

This work was supported by the Department of Defense grants W81XWH-16-1-0211 and W81XWH-16-1-0071 (A.F.R., A.A.P), Department of Veterans Affairs Merit award 1I0RX002101 (A.F.R., A.A.P.), and Department of Veterans Affairs Career Development Award IK2 RX002010 012020 (L.P.S, A.F.R). This work was also funded by a Pilot project awarded under The Center for the Prevention and Treatment of Visual Loss (award C9251-C) from the Department of Veterans Affairs Rehabilitation R&D Services (A-S.W.). A-S.W. was also supported for this work by the International Headache Academy research award from the American Headache Society. A.A.P. was supported for this work by the Brockman Foundation, Elizabeth Ring Mather & William Gwinn Mather Fund, S. Livingston Samuel Mather Trust, G.R. Lincoln Family Foundation, and the Louis Stokes VA Medical Center resources and facilities. The contents do not represent the views of VA or the United States Government.

